# Predicting metabolic adaptation from networks of mutational paths

**DOI:** 10.1101/115170

**Authors:** Christos Josephides, Peter S. Swain

## Abstract

Competition for substrates is a ubiquitous selection pressure faced by microbes, yet intracellular trade-offs can prevent cells from metabolizing every type of available substrate. Adaptive evolution is constrained by these trade-offs, but their consequences for the repeatability and predictability of evolution are unclear. Here we develop an eco-evolutionary model with a metabolic trade-off to generate networks of mutational paths in microbial communities and show that these networks have descriptive and predictive information about the evolving communities. We find that long-term outcomes, including community collapse, diversity, and cycling, have characteristic evolutionary dynamics that determine the entropy, or repeatability, of mutational paths. Although reliable prediction of evolutionary outcomes from environmental conditions is difficult, graph-theoretic properties of the mutational networks enable accurate prediction even from incomplete observations. In conclusion, we present a novel methodology for analyzing adaptive evolution and report that the dynamics of adaptation are a key variable for predictive success.

---

Elucidating the factors that influence the emergence, diversity, and stability of microbial communities is a central interest in both ecology and evolution [1]. To predict and control community structure and function, it is necessary to understand how interactions between microbes and the environment manifest as selective pressures driving microbial adaptation and diversification [2].

Microbes frequently grow on mixtures of metabolic resources where competition for growth-limiting substrates is a ubiquitous selection pressure. Surveys have revealed that microbes do not simultaneously use all available substrates; instead, each species in a community specializes to a few substrates [3, 4]. This observation is anticipated by ecological (resource-ratio) theory, which posits that constraints in the capacity of an organism to use multiple substrates are necessary for diversifying selection in homogeneous environments [5], and is supported by competition experiments where trade-offs in using one substrate over another maintain metabolic polymorphisms [6, 7, 8, 9, 10].

The conditions under which a microbial population can invade another, and whether a stable community can be formed, are the subject of ecological invasion analysis. These analyses typically either assume competition between infinitesimally-varying phenotypes [11, 12] or are not concerned with the mutational paths [13, 14] that may be generated through successive mutations and invasions. Advances in experimental evolution, however, now enable tracking of microbial lineages for hundreds of generations [15, 16] and expose how sequences of mutations shape the evolutionary process through competition between possibly disparate phenotypes [8]. Moreover, these experiments reveal evolutionary trajectories with both parallel and unique dynamics [17, 18], as well as variability in long-term outcomes [10].

To investigate how the interplay between metabolic constraints, environmental conditions, and the distribution of mutations influences the adaptation process, we developed a model that combines microbial chemostat ecology with an evolutionary process. At the ecological level, microbes compete for two growth-limiting substrates, and we incorporate a phenomenological metabolic trade-off between the consumption of one substrate relative to the other. Cells inherit this phenotype through nearly-faithful clonal reproduction, but rare mutations can change the degree of substrate specialization. We do not restrict the size-effect of mutations so that, in the extreme case, any phenotype can mutate to any other. We determine and analyze all mutational paths and stable microbial communities that may arise to describe the entire adaptation process.

We begin by introducing the eco-evolutionary model and show that adaptation dynamics over mutational paths can be visualized as a network through constructing a Markov process over a finite set of possible communities. Multiple long-term behaviors emerge, including quasi-periodic outcomes and outcomes with many exclusive stationary states. Next, we show that these evolutionary outcomes are not restricted to distinct environments and that small environmental perturbations can change one type of outcome to another. In spite of this sensitivity to environmental conditions, the processes that lead to each evolutionary outcome have characteristic adaptation dynamics, which are reflected in graph-theoretic properties of the mutational paths. Finally, we show that evolutionary outcomes can be predicted using these network properties from incomplete observations of evolving microbial communities — even without knowledge of environmental conditions.

## Results

### Microbial ecology in the chemostat with two substrates

Microbial evolution is often studied in chemostats, and we focus on a chemostat model of microbial ecology [19, 20]. To investigate metabolic adaptation on multiple substrates, we extend the standard chemostat model [21] to include two growth-limiting substrates, which we name *u* and *v* (Supplementary Section 1.1). The two substrates are individually sufficient for microbial growth — i.e. they are perfectly substitutable [22]. Both substrates are added continuously to the chemostat, and cells grow in proportion to their concentrations. The chemostat is diluted at a constant rate, and populations must therefore grow at least as fast as the dilution rate to survive being washed out. At steady-state, the growth rate of a surviving population is equal to the chemostat dilution rate (Fig. 1A).

**Figure 1:**
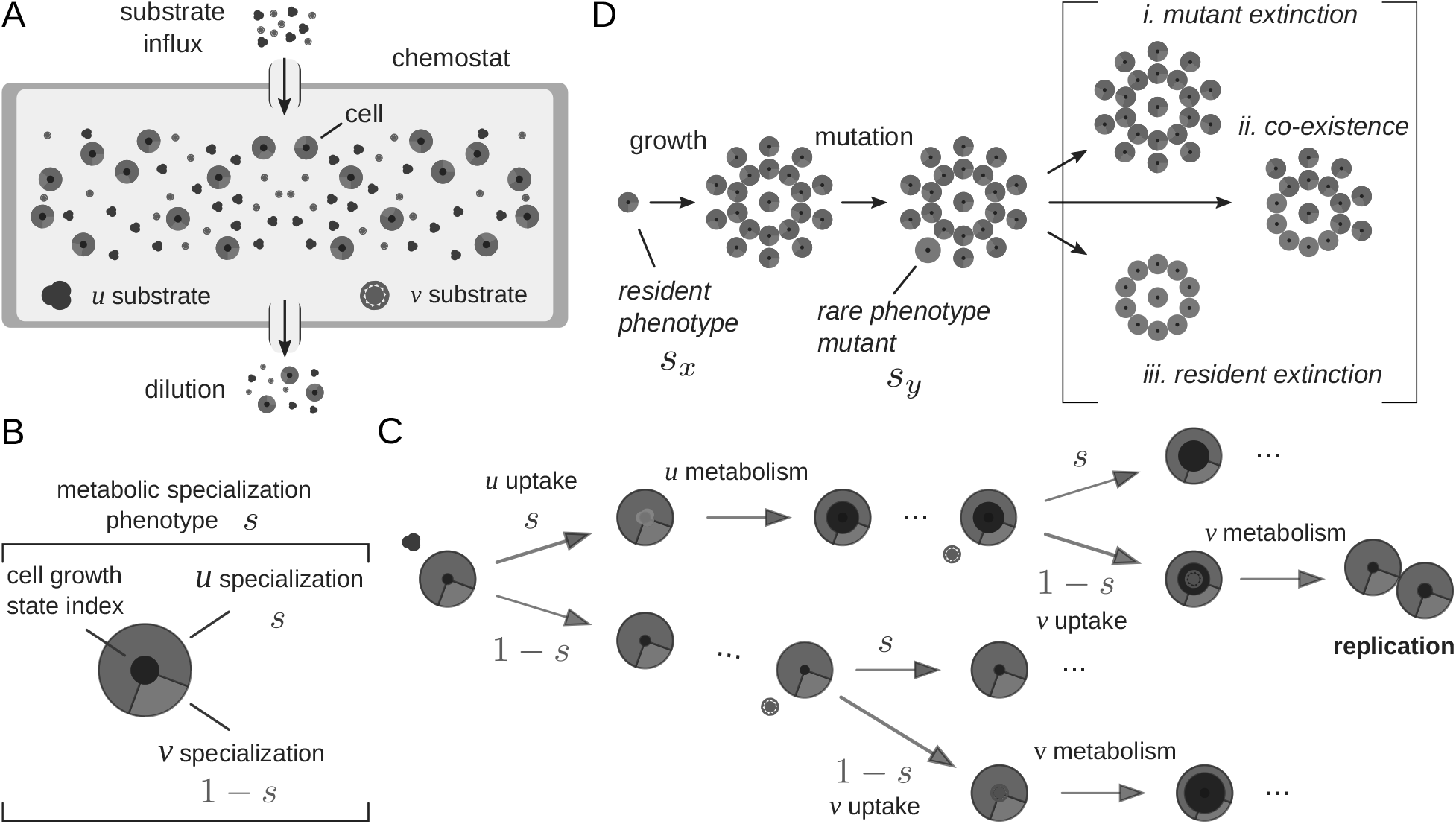
We model microbial ecology in the chemostat with two growth-limiting, substitutable, substrates (*u* and *v*) and a metabolic trade-off where specialization in one substrate reduces specialization in the other. **A**. The chemostat is a homogeneous environment with continuous addition of the two substrates and continuous dilution at fixed rates. The model is parameterized by the chemostat’s dilution rate, and a pair of influx rates, import rates, metabolic rates, and yields for the two substrates. **B**. The metabolic specialization of a cell is parameterized by a number between zero and one, *s*: values near zero indicate a *v*-specialist; values near one indicate *u*-specialist; intermediate values are substrate generalists. **C**. Cells increment their growth state by different amounts depending on the yield of the substrate. Cells replicate when their growth state exceeds a critical value. When a cell encounters a *u* substrate molecule, the probability that it will successful import this molecule is its phenotype value (*s*); when a cell encounters a *v* substrate molecule, the probability that a cell imports a molecule of *v* substrate is (1 − *s*). D. Competition between a (high abundance) resident phenotype and a (low abundance) invading phenotype follows a rare mutation event and has three possible qualitative outcomes.

Cells in the chemostat encounter both *u* and *v* substrates but cannot simultaneously specialize to using both. To investigate how a metabolic constraint affects adaptation [23], we consider a phenomenological trade-off between the import capacity for the two substrates (Supplementary Section 1.2). We parameterize the metabolic specialization of each cell by a number between zero and one, *s*: values near zero indicate that the cell specializes to *v*; values near one indicate *u* specialization. Cells inherit their phenotype (their value of *s*) from their parent nearly always without variation through clonal replication and cells cannot change their phenotype (Fig. 1B).

We can interpret the metabolic specialization, *s*, in two ways, using a common mathematical formulation (Supplementary Section 1.5). In the first interpretation, cells can always import both substrates but increase the rate of import for *u* at the expense of decreasing the import rate for *v*. The maximal import rate for *u* is multiplied by *s*; the maximal import rate for substrate *v* is multiplied by (1 − *s*) (Supplementary Section 1.3). Such a constraint may arise if a finite resource is shared between the production of the *u* and *v* permeases [24, 25, 26] or through antagonistic pleiotropy [27, 6, 7]. In the second interpretation, cells adopt a mixed strategy (following evolutionary game theory [28]) of randomly switching between two metabolic programmes, each exclusive to only one substrate. Then, the probability of adopting the *u* metabolic programme is *s* and the probability of adopting the *v* programme is (1 – *s*) (Supplementary Section 1.4).

We model the microbial life cycle through a sequence of substrate import, metabolism, and growth processes. Specifically, we structure the microbial population to discrete growth states that the cell progresses through to replicate. Cells metabolize *u* and *v* molecules at substrate-specific rates and obtain a substrate-specific yield in terms of growth state increments. Cells replicate when they have accumulated enough yield to exceed a threshold number of growth states (Fig. 1C).

A cell will almost always produce offspring that inherit its metabolic phenotype, but, rarely, a mutation will result in a phenotypically-distinct population. The new mutant will compete with the pre-existing resident population for extracellular substrates. We follow the theory of adaptive dynamics and include phenotype mutations only on evolutionary timescales [11, 12]: we assume that mutations are sufficiently rare that mutant phenotypes emerge only after the chemostat model has reached steady state. The ecological and evolutionary timescales are separated in this weak-mutation limit.

In our spatially homogeneous model at most two populations can co-exist on two substrates because of the competitive exclusion principle [29]. A mutant population is at first rare and must grow faster than the dilution rate to survive wash-out in the chemostat. When the chemostat contains a single phenotypic population, for example, invasion by a mutant has three possible outcomes: the mutant becomes extinct, the resident becomes extinct, or the two phenotypes co-exist in a community (Fig. 1D).

### Markov process of mutation-limited adaptation

Frequency-dependent selection gives rise to multiple mutational paths, which are contingent on the sequence and outcome of mutation and invasion events. Rare mutations generate a series of competition events between the resident and mutant populations with different metabolic phenotypes. Successful invasion by a new population modifies the chemostat environment through changing the steady-state levels of the substrates, and therefore the context in which future mutation and invasion events occur is modified through such ecological niche construction and destruction [23, 8, 2, 30, 18].

To generate the mutational paths, we first create an invasion map for the outcome of competitions between dynamically-stable communities of resident phenotypes and possible mutant phenotypes that may invade. To do so efficiently, we developed a dynamic programming algorithm to simulate invasion events assuming rare mutations on a discretized phenotype space (Fig. 2A). Briefly, the algorithm treats the invasion map as a tree: nodes are communities of phenotypes that can co-exist at steady-state and these are connected by edges representing single mutation and invasion events. The algorithm constructs the tree by iteratively perturbing each steady-state (parent node) to introduce a small mutant population. The resulting competition is simulated (Supplementary Section 1.6), and the steady-state outcome is analyzed and recorded as a connected steady-state (child node). We avoid redundant simulations, and so avoid recursive expansions of the tree, by appropriately terminating branches (Supplementary Section 2.1).

**Figure 2:**
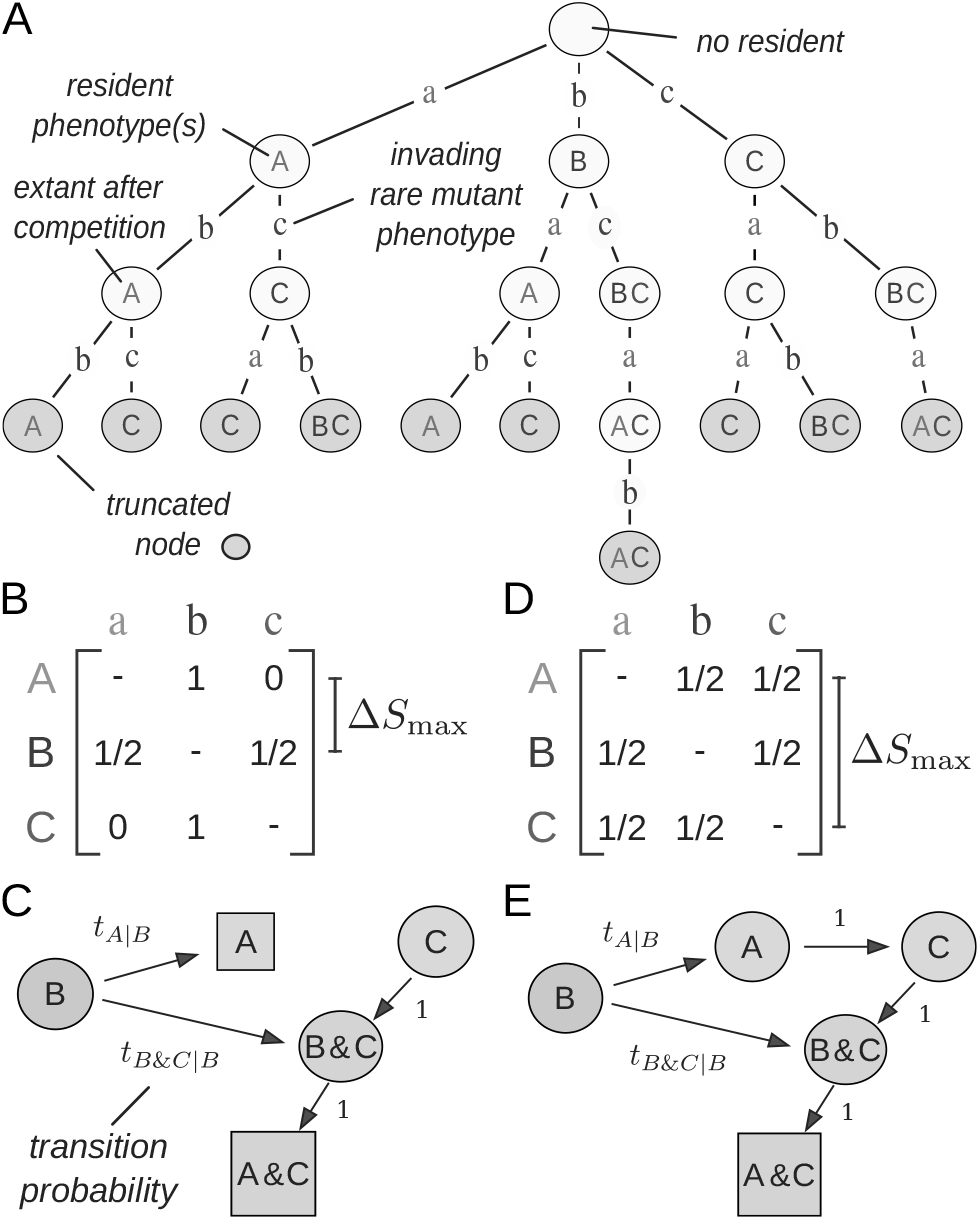
The Markov process of adaptation is generated from two components — simulations of ecological invasion and calculations of the transition probabilities based on the probabilities of each potential mutation — and is demonstrated with an example that has three phenotypes (labeled *A,B,* and *C*). **A**. To discover all dynamically-stable populations and communities, we simulate mutation and invasion events in a space of discrete phenotypes assuming rare mutations. We construct an invasion map as a tree: nodes are steady-states labeled with extant phenotypes. Each state is perturbed with a small mutant population and the resulting steady-state after competition is connected to the parent node. The algorithm dynamically truncates the tree to avoid any recursion. **B**. To calculate transition probabilities between resident states, we parse the invasion map in *A* with a mutational process. We assume uniformly distributed mutations with a maximum mutation size, ∆*S*_max_, centered on the resident phenotype. The matrix elements show the conditional probability that phenotype *s_x_* generates mutant phenotype *s_y_*. **C**. The Markov process in *B* visualized through its directed network graph. States are classified as either transient (circles) or recurrent (squares). **D**-**E**. As in *B-C,* but with a larger maximum mutation size (∆*S*_max_). The recurrent states, as well as the mutational paths of the process, depend on the distribution of mutations.

To define a Markov process of adaptation, we require the probability that a community will transition from one state to another as a result of a single mutation and invasion event (Supplementary Section 2.2). The probability of such a transition is proportional to the abundance of cells in the source state and to the propensity with which each cell generates the mutation that effects the transition to the target state (according to the deterministic invasion map). We will only be concerned with the sequence of transitions between one community state to another and not with the waiting time between such events. In this case, the distribution of mutations is the most important factor for determining which of the possible states will next follow a current state.

We use a uniform distribution of mutations in phenotype space centered on a resident phenotype, *s_x_,* with maximum mutation size ∆*S*_max_. Mathematically, a mutant phenotype *s_y_* can be generated by a cell with phenotype *s_x_* if |*s_y_* − *s_x_* | ≤ ∆*S*_max_. The mutation probabilities are chosen so that all of the discrete *s_y_* values that can be generated from *s_x_* are equally likely after correcting for boundary effects (Fig. 2B,D). The distribution of mutations affects both the mutational paths and the stationary behavior of the adaptation process, and we parse the invasion map using uniform distributions with different maximum mutation sizes to investigate this dependency.

To complete the description of mutation-limited adaptation as a Markov process, we must choose an initial distribution of ancestral states. Most laboratory evolution experiments start with isogenic strains, and, similarly in our model, adaptation starts with equal probability from any viable population comprising a single phenotype (rather than a community).

To determine the long-term evolutionary outcome of an adaptation process, we classify the states of the Markov process as either transient or recurrent. Transient states represent microbial communities that may only be visited once on a mutational path. A recurrent state, however, will always re-emerge once established, and the endpoints of mutational paths are always recurrent states (Fig. 2C,E). Recurrence occurs either when a microbial community cannot be invaded by any mutant that can be generated from the community (an evolutionarily-stable state [28]) or if there is a sequence of mutation and invasion events that returns to the community.

### Hierarchy of evolutionary outcomes

To investigate the effect of the environment and mutational parameters on the adaptation process, we randomly sampled 10, 000 environmental parameter sets (Supplementary Section 3.1). For each set of parameters, we calculated the invasion map using 11 discrete phenotype values ranging from a pure *v*-specialist to a pure *u*-specialist: *s* ∈ {0.0, 0.1,…,0.9,1.0}. We parsed each invasion map with uniform mutation distributions with increasing maximum mutation sizes, ∆*S*_max_ ∈ {0.1, 0.2,…, 0.9,1.0}, to generate ten adaptation processes. We then analyzed the 100, 000 resulting adaptation processes to determine their long-term evolutionary behavior.

We classify the evolutionary outcome of an adaptation process according to the number and type of its recurrent states (Fig. 3A). At a high level, we differentiate between processes with either a single recurrent state or multiple recurrent states. A recurrent state may be a dimorphic community, with two-coexisting phenotypes, or a monomorphic population with a single phenotype. We classify phenotypes as either (*u* or *v*) substrate specialists, when *s* ∈ {0.0,1.0}, or substrate generalists, when *s* ∈ {0.1, 0.2,… 0.8, 0.9}, and we distinguish between the possible community combinations of specialists and generalists in a recurrent state.

**Figure 3:**
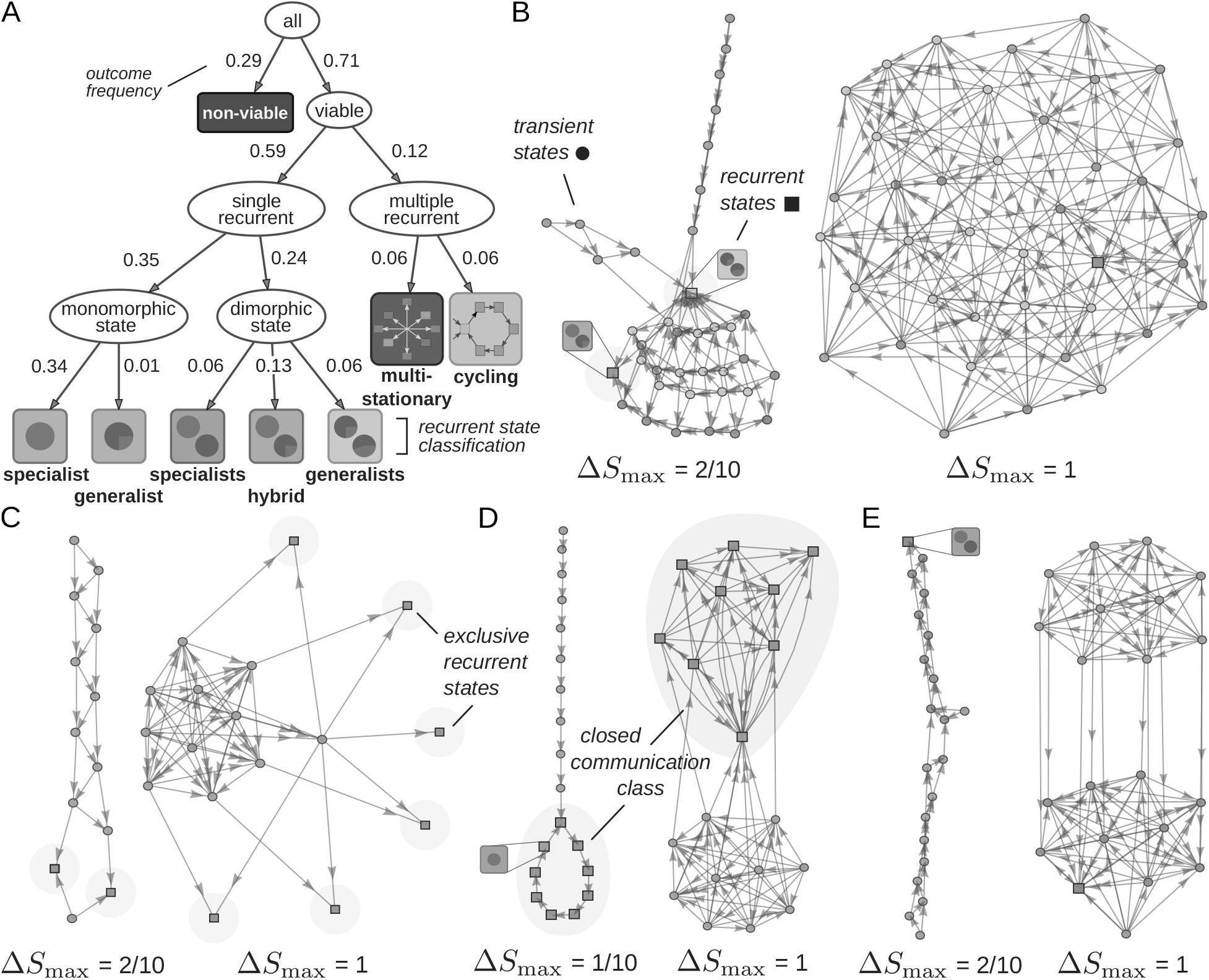
The model generates multiple evolutionary outcomes and complex networks of mutational paths. Qualitative network properties of adaptation dynamics can be visualized and conveyed through graphical representation. **A**. We hierarchically classify adaptation processes according to their evolutionary outcome using the number and type of recurrent states. Labels show the outcome frequencies obtained from parameter sampling. **B**-**E**. Networks of mutational paths from four sets of environmental parameters demonstrate the scope of the model’s dynamic and stationary behavior. Circles are transient states; squares are recurrent states; colors denote the number and type of metabolic phenotypes in each (microbial community) state following *A*. **B.** The maximum mutation size (∆*S*_max_) affects the mutational paths and long-term outcomes of adaptation (compare recurrent states between *left* and *right* networks). **C.** A process with multiple, non-connected, recurrent states has more than one evolutionarily stable state, each of which is reached with some (stationary) probability. **D.** A process where multiple recurrent states are connected exhibits quasi-periodic evolutionary cycling. **E**. An example of a potentially bottle-necked process. The network consists of two highly-connected (top and bottom) subgraphs, which are themselves connected via only a few mutation and invasion transitions.

We observed two qualitatively different behaviors in processes with multiple recurrent states, which we named ‘multi-stationary’ and ‘cycling’. Multi-stationary outcomes have many exclusive recurrent states where each recurrent state is reached with some probability. We found that the number of recurrent states and their stationary probability depended on the maximum mutation size (Fig. 3B, C). In these cases, we expect replicate evolution experiments to eventually show divergent phenotypic variability [8, 10, 31]. Cycling outcomes, on the other hand, are processes where no microbial community is completely resistant to invasion, and the adaptation process continually cycles between a set of states through mutation and invasion events. This cycling can be either periodic (Fig. 3D *left*) or aperiodic and unpredictable — albeit confined to a closed class of recurrent states (Fig. 3D *right).* These community changes occur at the evolutionary time-scale [32]. For example, in a dimorphic community of one specialist and one generalist population, mutation and invasion events can drive the generalist to extinction; this extinction leaves an unexploited metabolic niche that a newly-emergent generalist population can fill, thereby repeating the cycle (Fig.3D). Both of these evolutionary outcomes are qualitatively reproduced in stochastic simulations of mutational paths on a continuous phenotype space (Supplementary Section 2.3).

Visualizing the networks not only conveys information on the evolutionary outcomes but also on the adaptation dynamics. For example, we found that some adaptation processes had bottle-necked mutational paths: all mutational paths have to transition through a few common community states before reaching a recurrent state (Fig. 3E). We will later quantify these network properties to characterize the adaptation dynamics and construct a predictive model.

### Sensitivity of adaptation to environmental conditions

In light of the multiplicity and complexity of evolutionary outcomes, we investigated how particular outcomes are associated with the environmental parameters of the model. These parameters describe the chemostat set-up (substrate influx and dilution rates) and the properties of the two substrates (their yields, maximal import rates, and metabolic rates).

Although some evolutionary outcomes were more likely to be associated with certain environments, no clear pattern emerged (Fig. 4A). Using a nearest-neighbors algorithm to estimate the density of evolutionary outcomes, we observed that metabolically diverse (dimorphic) communities are most likely to emerge when the influx rates for the two substrates are similar and when the dilution rate is low. As the disparity in supply of the substrates increases, we find that communities undergo evolutionary cycling, perhaps signalling the onset of ecological collapse [33]. At even greater ratios of substrate influx, microbial communities disappear, leaving a population of a single specialist as the only evolutionary outcome.

**Figure 4:**
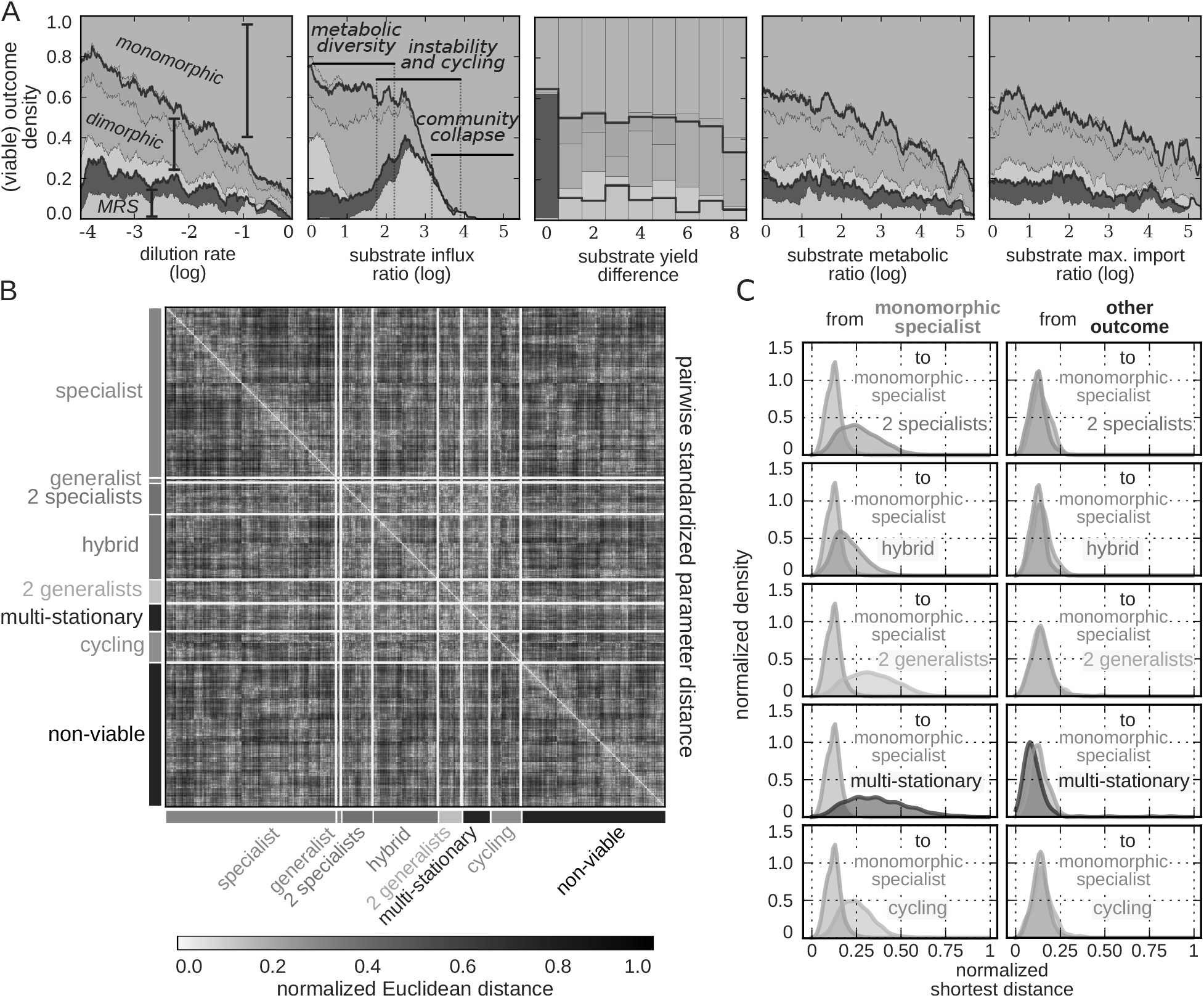
Complex association between the environment and evolutionary outcomes undermines the stability of microbial communities and complicates prediction. **A**. Evolutionary outcomes are more likely to be associated with some parameter values, but without a simple dependence. We estimated the empirical outcome densities from parameter sampling using a nearest-neighbors algorithm. For all parameter combinations except the dilution rate, we show densities at absolute value because of the symmetry about zero. **B**. Evolutionary outcomes form clusters in parameter space but these are not distinct. We used Euclidean distance in a standardized parameter space to perform within-outcome hierarchical clustering and then aligned clusters with samples from other outcomes to assess their overlap. We only plot a representative sample (1%) of the dataset, which preserves the observed frequencies of each outcome. **C**. Environmental perturbations often cause community collapse and only rarely create diverse communities. We calculated the standardized parameter distance from each process to the closest process in all other outcome classes to form the distribution of shortest-distances. Monomorphic specialists are generally closer to other monomorphic specialists than they are to outcomes with diverse communities *(left*). In contrast, processes with diverse outcomes are as close to each other as they are to monomorphic specialists *(right*). Here ‘other outcome’ refers to the labeled outcome in each panel that is not a monomorphic specialist.

Even by using combinations of environmental parameters to characterize outcomes, substantial unpredictability remained. We developed a hierarchical classification model, which combines six statistical estimators to make predictions (Supplementary Section 3.2). Its overall performance (a mean recall of 0.78 over the eight evolutionary outcomes), however, did not suggest that a reliable predictive map from environmental parameters to evolutionary outcomes could easily be formed.

We can partly understand this unpredictability by considering how the different evolutionary outcomes are distributed in parameter space, where distinct clusters do not appear to exist. For each outcome, we first performed hierarchical clustering using the pairwise Euclidean distance in a standardized parameter space (Supplementary Section 3.3) and then aligned these clusters with samples from other outcomes to assess their overlap (Fig. 4B). In general, while parameter sets that produced the same outcomes did form clusters, these were not distinct: a sample or cluster from one outcome is typically proximal to samples or clusters from other outcomes. Environmental perturbations are therefore often likely to cause a qualitative shift in the long-term community.

In particular, monomorphic specialists ‘permeate’ parameter space, and are typically close to all other outcomes. By calculating the distribution of shortest-distances from one type of outcome to either the same outcome or one of the other seven possibles outcomes, we can estimate the magnitude of the environmental change required to both maintain the type of outcome and to alter one outcome to another (Supplementary Section 3.4). Assuming that the probability of environmental perturbation is inversely proportional to its magnitude, a change in the environment is approximately as likely to preserve a diverse community as to lead to its collapse to a monomorphic specialist (Fig. 4C, *right*). The converse, however, is not true: a monomorphic specialist population is far less likely to transition to a diverse community (Fig. 4C, *left*).

We conclude that prediction of evolutionary outcomes from environmental conditions, even in simple chemostat environments, is likely to be challenging. Evolutionary outcomes are only the endpoints of the adaptive process, however, and we next investigate if the dynamics of these processes (the mutational paths) are characteristic of their endpoints.

### Adaptation dynamics and mutational paths

The mutational paths of the Markov process describe the dynamics of adaptation, and the processes we construct have tens to millions of mutational paths in spite of a relatively small number (eleven) of discrete phenotypes and a maximum of two co-existing populations in each community.

We analyze adaptation dynamics by enumerating the mutational paths of a process (Supplementary Section 4.1), calculating their probabilities, and determining their properties (Fig. 5A). For example, we define the path length as the number of state transitions, from an ancestral state to a recurrent state, in one realization of the adaptation process (a single run of an evolution experiment).

**Figure 5:**
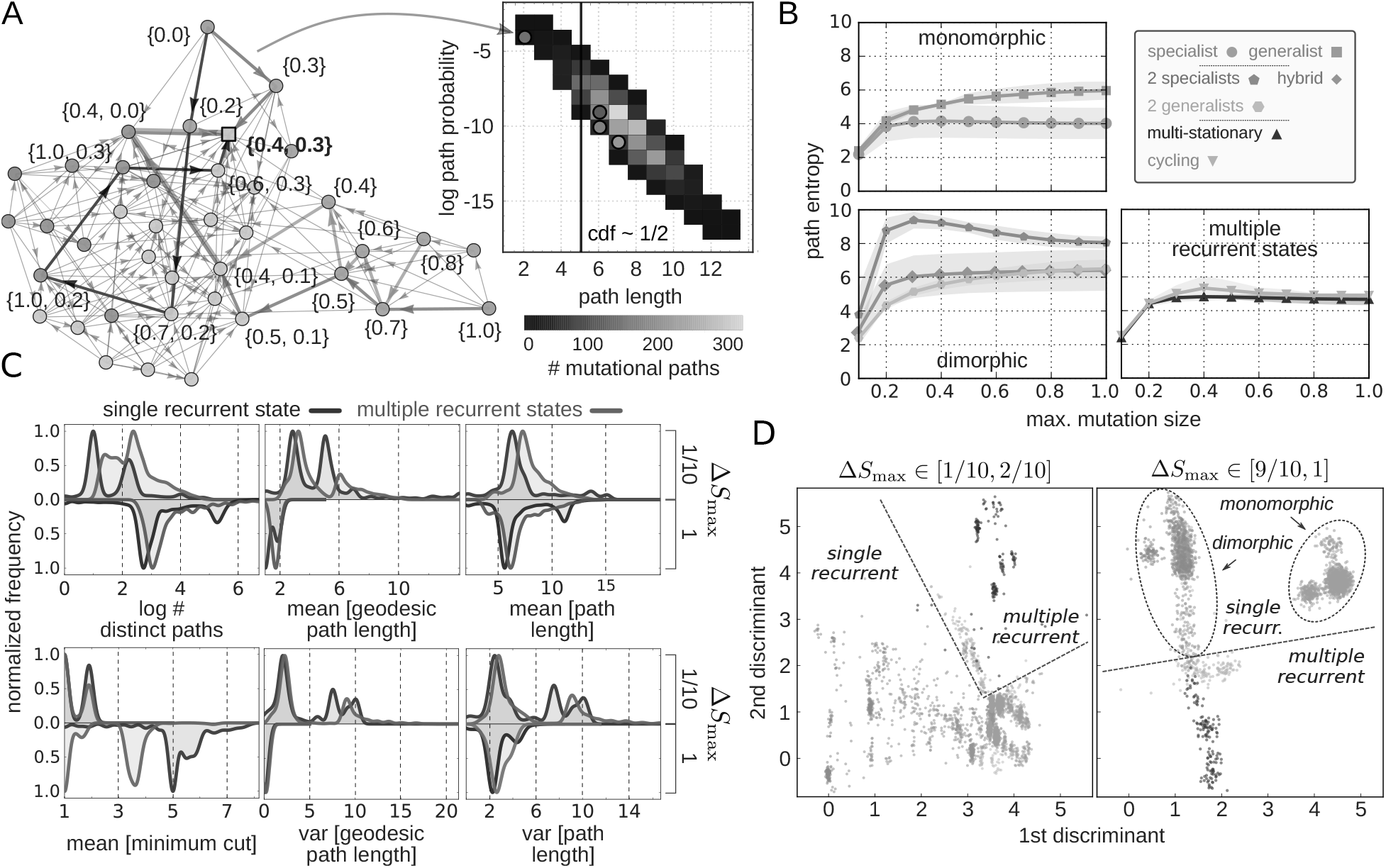
Processes with different evolutionary outcomes have characteristic adaptation dynamics, which can be quantified and compared through the properties of their mutational paths. **A**. We enumerate mutational paths to generate the path distribution. Paths begin at a monomorphic state (with equal probability) and end at a recurrent state. The path probability decreases in proportion to the length of the path. Most mutational paths in a network are of intermediate length, and only a few paths are either short or long. Statistical properties of adaptation, like the mean and variance of the path length, are calculated from the path distribution. **B**. The repeatability of adaptation dynamics depends on the long-term evolutionary outcome and typically increases with maximum mutation size. The entropy of the path distribution quantifies repeatability (high entropy implies low repeatability). Points show the mean path entropy for networks with different outcomes; shaded regions are ± the standard deviation. **C**. Mutational path statistics vary with the maximum mutation size and the process’s evolutionary outcome. We plot the distributions for six statistical path properties, grouping by the number of recurrent states and two extremes of maximum mutation size. The distributions are multi-modal, and the peaks correspond to outcomes lower in the classification hierarchy (Fig. 3A). **D**. Processes with different evolutionary outcomes have characteristic adaptation dynamics. Linear combinations of the six path properties in *C* that best separate the adaptation processes in a twodimensional projection were identified through linear discriminant analysis. Lines and ellipses were drawn manually.

The degree to which adaptive evolution is repeatable is of long-standing interest [34, 35], but analyses are typically restricted to models with static fitness landscapes [36]. To quantify the repeatability of adaptation in our model, where fitness landscapes change dynamically through eco-evolutionary feedbacks, we mathematically define repeatability as the entropy of the path distribution. If replicate experiments are likely to follow only a few mutational paths, the entropy is small and repeatability is high; if each replicate experiment follows a different mutational path, the entropy is large and repeatability is low.

In our model, the repeatability of adaptation dynamics is associated with the evolutionary outcome of the adaptation process and varies with the maximum mutation size (Fig. 5B). Path entropy typically increases with maximum mutation size as more mutational paths become possible. Notably, however, dimorphic specialist outcomes have a non-monotone path entropy with maximum mutation size (Fig. 5B, blue line) because a few mutational paths with large mutations come to dominate the process. We find that processes where monomorphic specialist populations are evolutionarily stable have the most repeatable dynamics; in contrast, processes with metabolically diverse communities have the least repeatable adaptation dynamics.

To compare the dynamics between adaptation processes, we constructed a condensed dynamical profile for each through calculating six statistical properties of its mutational paths. These properties were: the number of paths, the mean and variance of geodesic (shortest) path length, the mean and variance of all paths, and the mean minimum cut size (the smallest number of edges that must be removed from the graph to disconnect an initial state from a recurrent state — a measure of the extent of bottle-necks in the process).

We analyzed the six mutational path properties for processes with different evolutionary outcomes and at different maximum mutation sizes (Fig. 5C). The effect of increasing the maximum mutation size depends on the evolutionary outcome of the process, but the qualitative trends are consistent. Specifically, larger mutations decrease both the mean and the variance of the geodesic path length because recurrent states can be reached via fewer mutation and invasion events of larger effect. Larger mutations have a similarly decreasing, albeit smaller, effect on the mean and variance of the overall path length. Finally, we found that a larger maximum mutation size generally leads to processes with fewer bottle-necks (higher minimum cut size), particularly for processes with a single recurrent state.

The distributions of mutational path properties are multi-modal (Fig. 5C), where the peaks correspond to evolutionary outcomes low in the outcome hierarchy (Fig. 3A), suggesting that processes with different evolutionary outcomes have characteristic adaptation dynamics. We used discriminant analysis to find linear combinations of the six path properties that maximize the separation of the seven viable evolutionary outcomes according to their dynamics. We found that different outcomes, particularly those higher in the outcome hierarchy (Fig. 3A), occupy different regions in a two-dimensional projection of the discriminant space (Fig. 5D). We can therefore conclude that mutational paths characteristically identify processes’ longterm outcomes, and we contrast this positive finding with the difficulty of obtaining an association between environmental parameters and evolutionary outcomes (Fig. 4).

The path properties we calculate quantify statistical aspects of the adaptation process; however, enumeration of paths becomes computationally prohibitive for larger networks and also relies on correct classification of states. We next present an alternative graph-theoretic approach that relaxes these constraints and forms the basis of a model for predicting evolutionary outcomes from observations of adaptation dynamics.

### Predicting evolutionary outcomes from networks of mutational paths

To motivate our predictive model, we consider an evolution experiment as the progressive construction of a network of mutational paths. In such an experiment, the resident microbial populations in a chemostat [37] are periodically assayed, and the transitions between resident communities are used to construct the network. To simulate such a procedure, we re-sampled the 100, 000 adaptation processes in our dataset to obtain networks at varying stages of completion (Supplementary Section 5.1). Incomplete networks may contain spurious recurrent states, which obfuscate the process’s evolutionary outcome (Fig. 6A, *top*). These states appear recurrent because the mutation and invasion events that lead away from these states have not yet been observed in the experiment. The predictive objective, then, is to forecast the true recurrent states of the complete network from an incomplete sample.

**Figure 6:**
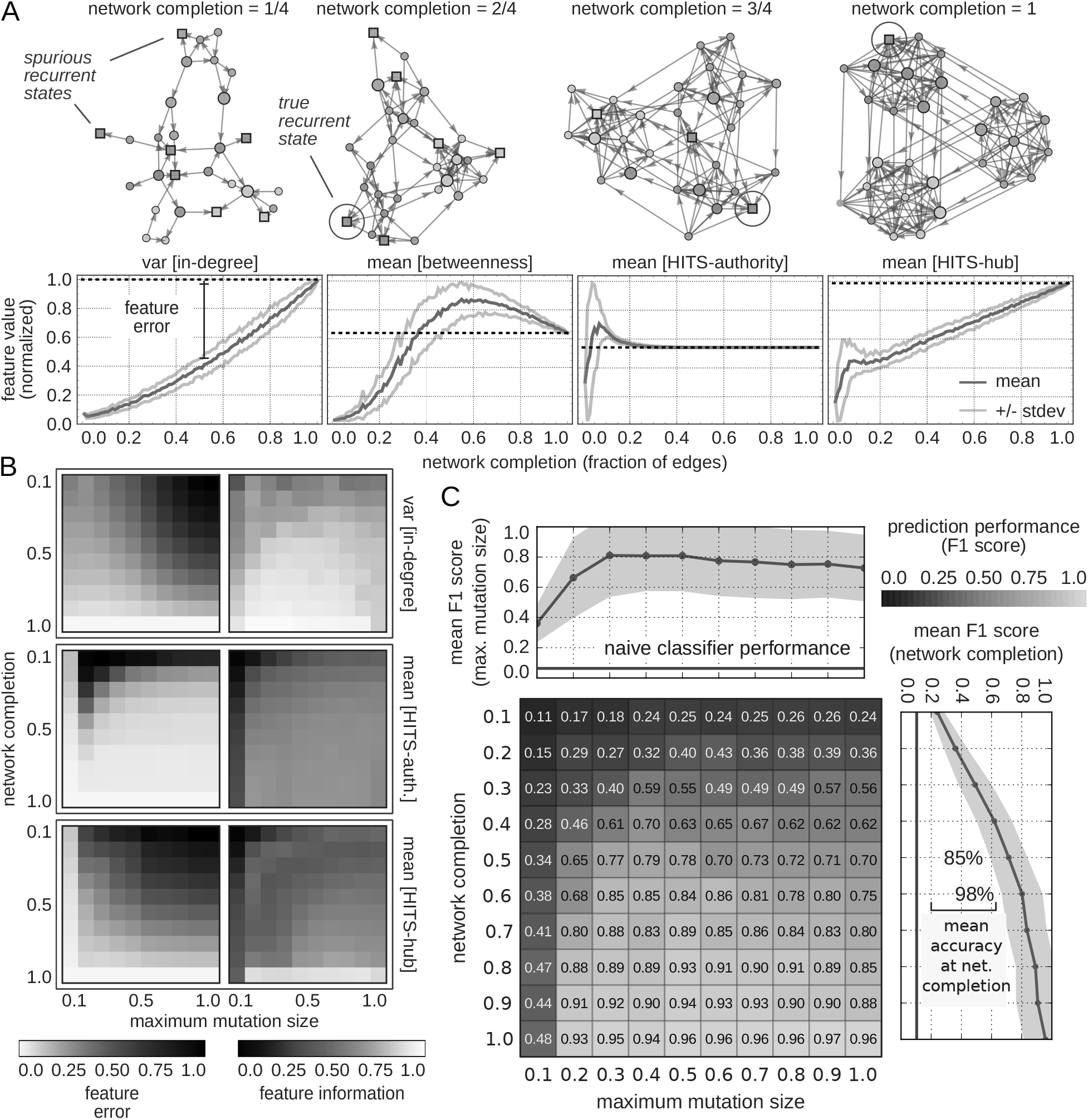
Evolutionary outcomes can be predicted from graph-theoretic properties of incomplete networks of mutational paths. To generate incomplete networks we sampled networks to varying degrees of completion. For both complete and incomplete networks, we calculated and summarized six network centralities, which characterize different aspects of network topology, through their mean and variance. Statistics for the twelve centralities summarize mutational networks through a low-dimensional space. **A**. The state of network completion affects its topological properties, which is reflected in the centrality statistics. For an example process (*top*), we show the progression of four centrality statistics (*bottom*) to highlight the variability in their convergence as a function of network completion. Dark blue lines show the mean and light blue lines show the standard deviation for 100 network samples at different stages of network completion. **B.** We characterized the centrality statistics both by how quickly they converge to their complete-network value (feature error, *left*) and by how well they can discriminate between evolutionary outcomes (feature information, *right*). The plots show the mean of feature error (the normalized difference between the complete- and incomplete- values of the centrality statistic) and the mean of the mutual information between the centrality statistic and the evolutionary outcomes. We found both invariant (e.g. mean of HITS authority) and informative (e.g. variance of in-degree) centrality statistics (Supplementary Section 5.2) as well as a dependency on the maximum mutation size. C. Evolutionary outcomes can be predicted with a statistical classifier trained on the twelve centrality statistics from partial and complete networks. We assessed classifier performance via the unweighted mean of the *F1* score, taken over the seven evolutionary outcomes, and report the mean performance on the test set for a ten-fold cross-validation. *Top* and *right* line plots show mean performance, with standard deviation in shaded regions, taken over the columns (network completion) and rows (maximum mutation size). The red line is the benchmark performance of a naive classifier, which predicts following the empirical frequencies of the outcomes.

Our predictive model relies on quantifying how networks change with accumulating observations. As we have shown earlier, evolutionary outcomes have characteristic mutational paths. To use the information encoded in the entire network of mutational paths at once, we measure graph-theoretic properties that quantify abstract features of the network. Specifically, for each network, we calculated six centrality measures [38] that characterize different aspects of network topology (Supplementary Section 4.2 and Supplementary Figure 6). Though less interpretable from an ecological or evolutionary perspective compared to path properties (such as path length) a network’s centrality distributions nevertheless contain valuable predictive information. For each network we calculate the six centrality distributions and summarize each through its mean and variance. As mutation and invasion events are observed, new vertices and edges are added to the network, and the resulting changes in network topology are reflected in changes in the centrality statistics (Fig. 6A, *bottom*).

We identified both invariant and informative topological changes as networks are progressively completed (Supplementary Section 5.2). We found that some centrality statistics converge to their complete-network value even when the network is mostly incomplete — remaining invariant as more vertices and edges are added — while others change considerably during network construction (Fig. 6B). Nevertheless, we noted that the most informative centrality statistics (those with distinct distributions for the different outcomes) converged gradually (Supplementary Figure 7).

Differences in network topology, even for incomplete networks, were sufficient to enable reliable prediction of evolutionary outcomes using a statistical-learning approach. Centrality statistics provide a projection of the mutational path network, through its topology, to a low-dimensional space. We trained a classifier to predict evolutionary outcomes from these centrality features using data from incomplete and complete networks, the degree of network completion, and the maximum mutation size (Supplementary Section 5.3). To avoid biasing against minority outcomes, we used the unweighted mean *F*1 score (over the outcome classes) as the optimization metric. Neither the degree of network completion or the maximum mutation size will typically be known during an experiment, and we marginalized over both these variables during testing (Supplementary Figure 8). The performance of the classifier improves with increasing network completion and is best at intermediate maximum mutation sizes (Fig. 6C). At 50% network completion, the classifier predicts the correct long-term outcome for about 85% of all adaptation processes (average *F*1 score > 0.7), which rises to 98% at 60% graph completion (*F*1 = 0.8).

We have shown here that our predictive approach can be useful in forecasting changes in the community, including the loss of (metabolic) diversity, through incomplete observations of adaptation dynamics. Detecting the onset of community transitions is an enduring problem, but most approaches focus on catastrophic transitions and analyze the dynamics of a single dynamical model [39, 40, 33]. Nevertheless, the loss of biodiversity following environmental disturbance can, as we have shown, involve a series of ecological transitions mediated by multiple mutation and invasion events, and our network approach can address this challenge.

## Discussion

We have shown that a simple model of metabolic adaptation comprising a trade-off in the use of one type of substrate over another can generate surprisingly rich behavior. On one hand, this behaviour is sufficiently complex that it is difficult to predict the outcomes of adaptation, even qualitatively, from environmental parameters. On the other, we have demonstrated that statistical properties of the mutational paths can uniquely characterize long-term outcomes. Thus, similarly to structural indicators in evolutionary game theory [41, 42] and statistical indicators in ecology [39, 40, 33], the dynamics of adaptation are a key variable for the successful prediction of community outcomes.

Our approach combines elements from the replicator-mutator equation [43, 44] and adaptive dynamics [11, 45, 12] to couple an ecological model with an evolutionary process. Ecological interactions between cells and the environment induce eco-evolutionary feedbacks [11, 45, 23, 2, 30, 18]: resident phenotypes define environmental niches for invasion by mutants, which in turn shape niches for future invasion. Such behavior is predicted to occur, for example, in the gastrointestinal tract [46, 47]. We formalized adaptation as a discrete-time Markov chain because the evolutionary model initiates ecological invasion via rare mutation events. This formulation emphasizes the dynamics of adaptation, i.e. the mutational paths [36, 34], rather than focusing on isolated phenotype competitions or evolutionary ‘end-points’.

Several extensions can be made, but are likely to only increase the complexity of the dynamics observed. For example, the substrates can be made partly substitutable [22, 5] or their number increased. Growth could be parameterized by several phenotypic traits combining multiple metabolic [48, 49] or cellular [26, 50] trade-offs. We ignored stochastic extinction of rare mutants, and we anticipate that such effects will change the statistical properties, but not the long-term outcomes, of adaptation. Finally, the model operates in the ‘strong-selection, weak-mutation’ regime [51], therefore we omit both multiple simultaneously invading mutants (clonal interference [52, 53]) and the occurrence of multiple mutations occurring within one invading lineage [54, 55].

The model demonstrates the importance of trade-offs for generating metabolic complexity in microbial communities [5, 1]. If a cell is able to evolve its response to one substrate unconstrained by its response to the other, then only one evolutionary outcome is possible — a phenotype that maximally depletes both substrates — because all other phenotypes will be unable to grow sufficiently fast in the chemostat to survive dilution [56, 21]. Metabolic trade-offs, however, preclude this single optimal phenotype and, together with dynamically changing environmental niches [8, 2, 18] and a limited distribution of possibly large-effect mutations [57], generate complex adaptation dynamics and long-term behaviors. Intracellular trade-offs are expected to be common [48, 26, 50], and our results support the idea that the resulting frustration of optimal behaviors is a major factor generating the complexity observed in microbial communities.

## Methods

### Model simulations

We modified the chemostat model [21] to include two substrates and structured populations (Supplementary Section 1.5), and simulated the model to steady-state using the numerical integration libraries in Mathemat-ica 10 [58] (Supplementary Section 1.5-1.6). Models were dynamically generated, by adding and removing invading and extinct populations, as required for the invasion map (Supplementary Section 2.1). The invasion map was parsed to calculate the state transition probabilities for the Markov process (Supplementary Section 2.2).

### Data analysis

We analyzed the adaptation processes from 10, 000 samples of environmental parameters (Supplementary Section 3.1). The path distribution, statistics of path properties (Supplementary Section 4.1), and centralities (Supplementary Section 4.2) were calculated for each network and are available as a tabulated dataset. Data was analyzed using the Python `scikit-learn`[59] library for statistical models. Incomplete networks were generated by re-sampling the complete-network dataset (Supplementary Section 5.1). Mutual information was calculated using non-parametric entropy estimation [60] (Supplementary Section 5.2). The predictive model was based on a Random Forest classifier (Supplementary Section 5.3).

## Acknowledgments

We thank Elco Bakker, Matteo Cavaliere, and especially Luke McNally for critical comments on the manuscript; we are grateful to Nikola Popovic for help with model analysis. C.J. acknowledges support from a Wellcome

## Author contributions

C.J. and P.S.S. designed research; C.J. performed research and analyzed data; C.J. and P.S.S. wrote the paper.

